# Metagenomic analysis reveals novel dietary-related viruses in the gut virome of marmosets hybrids (*Callithrix jacchus x Callithrix penicillata*), Brazil

**DOI:** 10.1101/2022.10.06.509726

**Authors:** Thamiris dos Santos Miranda, Francine Bittencourt Schiffler, Mirela D’arc, Filipe Romero Rebello Moreira, Matheus Augusto Calvano Cosentino, Amanda Coimbra, Ricardo Mouta, Gabriel Medeiros, Déa Luiza Girardi, Victor Wanderkoke, Caique Ferreira Amaral Soares, Talitha Mayumi Francisco, Malinda Dawn Henry, Bianca Cardozo Afonso, Flávio Landim Soffiati, Suelen Sanches Ferreira, Carlos Ramon Ruiz-Miranda, Marcelo Alves Soares, André Felipe Andrade dos Santos

## Abstract

Viral metagenomics has contributed enormously to the characterization of a wide range of viruses infecting animals of all phyla in the last decades. Among Neotropical primates, especially those free-living introduced, knowledge about viral diversity remains poorly studied. Therefore, through the use of metagenomics based on virus enrichment, we explored the viral microbiota present in the feces of introduced common marmosets (*Callithrix sp.*) in three locations from the Silva Jardim region in the State of Rio de Janeiro, Brazil. Fecal samples were collected from nine marmosets, pooled into three sample pools and sequenced on Illumina MiSeq platform. Sequence reads were analyzed using a viral metagenomic analysis pipeline and two novel insect viruses belonging to the *Parvoviridae* and *Baculoviridae* families were identified. The complete genome of a densovirus (*Parvoviridae* family) of 5,309 nucleotides (nt) was obtained. The NS1 and VP1 proteins share lower than 32% sequence identity with the corresponding proteins of known members of the subfamily *Densovirinae*. Phylogenetic analysis suggests that this virus represents a new genus, named *Tritonambidensovirus* due to telomeric structures at the 3’ and 5’ ends of the genome. The novel species received the name *Fecalis tritonambidensovirus 1*. The complete circular genome of a baculovirus of 107,191 nt was also obtained, showing 60.8% sequence identity with the most closely related member of the *Baculoviridae* family. Phylogenetic analysis suggests that this virus represents a new species of *Betabaculovirus*, named *Callithrix fecalis granulovirus*. In addition, sequences from several families of arthropods in the three pools evaluated were characterized (contigs ranging from 244 to 6,750 nt), corroborating the presence of possible insect hosts with which these new viruses may be associated. Our study expands the knowledge about two viral families known to infect insects, an important component of the marmosets’ diet. This identification in hosts’ feces samples demonstrates one of the many uses of this type of data and could serve as a basis for future research characterizing viruses in wildlife using noninvasive samples.

## 1. Introduction

Recent advances in high throughput sequencing (HTS) technologies have allowed metagenomic analysis to reveal an increasing genetic diversity of viral genomes ranging from unicellular bacteria, archaea and protozoa to multicellular organisms such as plants, fungi, invertebrates and vertebrates (Temmam *et al.*, 2014). Millions of wild fauna viruses are poorly characterized or totally unknown to man (Carroll *et al.*, 2018). Among vertebrates, little is known about the virome of Neotropical Primates (NP), especially among free-living populations. ADuarte *et al.,* (2019) characterized the fecal virome of free-living *Sapajus libidinosus* in the Brazilian Cerrado and identified novel known and putative viruses. However, itis currently estimated that NP are distributed in 183 species, 21 genera and three families (*Atelidae, Pitheciidae* and *Cebidae*), according to molecular data (Dumas and Mazzoleni, 2017; Perelman *et al.*, 2011; IUCN/SSC Primate Specialist Group, 2021), for which there are no systematic studies of viromes. Characterizing viruses infecting this group, as well as their susceptibility to infectious diseases, is key for understanding primates’ disease ecology, evolution and conservation.

The two species of common marmosets, the white-tufted (*Callithrix jacchus*) and the black-tufted (*Callithrix penicillata)* are diurnal, arboreal NP, native of northeastern and central Brazil, respectively (Malukiewicz *et al.*, 2021). Adults can weigh from 300 to 500 g and have an average life expectancy of 5 to 12 years in research colonies (Ross *et al.*, 2012). Marmosets live in family groups of up to 15 individuals with a dominant breeding pair (Digby, 1995). In the wild, their diet is mainly composed of plants, fruits, seeds, flowers, insects, and small animals (Pinheiro and Mendes Pontes, 2015). As a result of the illegal wildlife trade, these marmosets have been introduced to several regions outside and within Brazil, where they are considered a threat to endemic species (Pinheiro and Mendes Pontes, 2015). One of these regions is the Atlantic coastal forest of Rio de Janeiro state, Southeast Brazil. In this region, these marmosets occur as hybrids of both species, have been ecologically successful and are a threat to the entire population of endangered golden lion tamarins (*Leontopithecus rosalia*) (Malukiewicz *et al.*, 2015; Ruiz-Miranda *et al.*, 2006). The biology of marmosets is similar to lion tamarins, therefore they can readily compete for food, shelter and can transmit etiological agents not currently present in lion tamarins, becoming a concern for the conservation of the native species (dos Santos Sales *et al.*, 2010).

Analyses of the viral microbiota can be useful in efforts to preserve host biodiversity and to guide conservation strategies (Stumpf *et al.*, 2016). Feces might contain genetic material of pathogens, food sources (animals and/or plants), and commensal gut microbes, being an important resource for the analysis of ecological networks and the discovery of new viruses (Ge *et al.*, 2012; van den Brand *et al.*, 2012). In addition, feces are a non-invasive sample material, providing a safe alternative for biological sampling (Bergner *et al.*, 2019), since the small body size of many NP is a limiting factor for the collection of other types of samples, such as blood. Moreover, feces samples collection causes little to no animal stress, an ethically desirable feature.

Therefore, in this study, we characterized for the first time the viral metagenome of free-living introduced hybrids of *C. jacchus x C. penicillata* using stool samples. A potentially new genus and consequently a new species of Densovirus (family *Parvoviridae*) and a new species of *Betabaculovirus* (family *Baculoviridae*) were discovered through the analysis of HTS data, provisionally named *Fecalis tritonambidensovirus* 1 (FtDV) and *Callithrix fecalis granulovirus* (CafeGV), respectively. Evolutionary analyses indicate the new viruses are closely related to viruses previously characterized in insects, suggesting the reported experiment captured viral diversity associated with marmoset arthropod diet.

## 2. Material and Methods

### 2.1. Sample Collection

Fecal samples were collected from nine common marmosets living in three localities of Silva Jardim municipality, in the state of Rio de Janeiro, Brazil (**Table 1**). The marmosets were accustomed to regular human contact and consistently monitored by the *Associação Mico Leão Dourado* (AMLD). They were captured individually with Tomahawk^®^ live traps baited with bananas and placed on bamboo platforms positioned inside the forest fragment at 1.5 m above the ground. After capture, animals were transported to the laboratory of the AMLD, also in *Silva Jardim*, for sanitary evaluation and sample collection. In the laboratory, opportunistic fecal samples were collected on clean paper surfaces positioned under traps or directly collected from animals during manipulation. Animals were immobilized with injection of ketamine (approximately 10-15 mg/Kg of body weight) into the intramuscular region of the inner thigh and general information were collect for all sampled animals, such as: ID number, group, species, age, sex, weight, and clinical conditions. Each fecal sample was transferred to 15 mL falcon tubes with RNA*later*^™^ (Thermo Fisher Scientific, Walham, USA), a nucleic acid preservation buffer, with a volumetric ratio of feces-to-buffer of roughly 1:1. Samples were kept at room temperature and shipped to the *Laboratório de Diversidade e Doenças Virais* (LDDV) at *Universidade Federal do Rio de Janeiro* (UFRJ), Rio de Janeiro, Brazil, to be stored at −20 °C. All procedures were performed by veterinarians and biologists from AMLD and all samples were collected following the national guidelines and provisions of CONCEA (National Council for Animal Experimentation Control, Brazil), which included animal welfare standard operating procedures. This project was approved by the Ethics Committee on the Use of Animals (CEUA) of UFRJ (reference number 037/14). This study is reported in accordance to ARRIVE guidelines (https://arriveguidelines.org/resources/questionnaire).

**Table 1:** General sampling information.

| Species | Location | GPS | Sample Type | Library Code | Number of Individuals |
| --- | --- | --- | --- | --- | --- |
| *Callithrix sp.* | Afetiva, Silva Jardim, RJ | 22°37'58.84"S<br>42°28'1.47"W | Feces | S-Afe | 4 |
| *Callithrix sp.* | Igarapé, Silva Jardim, RJ | 22°30'27.41"S<br>42°18'34.66"W | Feces | S-Igar | 2 |
| *Callithrix sp.* | Iguape, Silva Jardim, RJ | 22°30'20.74"S<br>42°19'24.27"W | Feces | S-Igua | 3 |

### 2.2 High-Throughput Sequencing

To maximize sequencing efficiency, samples were pooled into three groups according to locality (Afetiva, Igarapé and Iguape) with four, two and three samples each, respectively (**Table 1**). The samples were vigorously vortexed until complete homogenization. Afterwards, approximately 1 mL of sample was transferred to the extraction bead tube (MP Biomedicals, Santa Ana, CA, USA) to disrupt the cells. The supernatant was collected by centrifugation at 6,000 × g for 10 min at 4°C, followed by filtering using Millex-HV 0.45 μm SLHV0135L (Merck & Co., Radington Township, NJ, USA) to remove host cells. The filtrates were submitted to an ultracentrifugation at 35,000 g for 90 minutes at 4°C to separate the enriched viral background fraction. The supernatant was discarded and approximately 0.2 mL of the sample was incubated with DNases (Promega and Epicentre, Madison, WI, USA), RNase A (Ambion, USA) and Benzonase (Sigma, USA) for 90 minutes at 37°C to digest unprotected nucleic acid. Viral nucleic acids were then isolated using QIAamp MinElute Virus Spin Kit (QIAGEN, Hilden, Germany) according to the manufacturer’s instructions with some modifications described *(Cosentino et al., 2022).* Complementary DNA (cDNA) synthesis was performed using the Superscript III First Strand Synthesis Supermix Kit (Thermo Fisher Scientific) and for the synthesis of the second strand of cDNA was used the Klenow fragment 3’-5’ exo (New England Biolabs Inc., Ipswich, MA, USA), following the manufacturer’s instructions. Mixed nucleic acid (DNA and cDNA) from all samples were subjected to library construction using Nextera XT DNA Sample Preparation Kit (Illumina Inc., San Diego, USA) and sequenced on an Illumina MiSeq platform (Illumina) at the Department of Genetics, UFRJ.

### 2.3 Bioinformatics Analyses

Sequencing data was processed with a custom pipeline. Raw reads were filtered with Fastp v.0.20.1 (Chen *et al.*, 2018), which removed adapters and low-quality reads (Phred score < 30 and fragment size < 50 bp). Contigs were *de novo* assembled with the Meta-spades software v.3.15.3 (Nurk *et al.*, 2017). Both reads and contigs from each sample were taxonomically assigned using Diamond v.2.0.14 (Buchfink *et al.*, 2015), with an e-value cut-off of 10^−5^ and parameters ‘--more-sensitive’ and ‘--max-target 1’ specified. The similarity search was performed against the complete non-redundant protein database from NCBI (as of July 27th, 2021). Interactive visualization plots were generated with Krona v.2.7.1 (Ondov *et al.*, 2011). Following (Farouni *et al.*, 2020), to avoid false positive results due to sequencing index-hoping (incorrect assignment of reads to a given sample), we considered as invalid the identification of a viral family when it was detected by a number of reads that was less than 1% of the highest count identified for the same family among all other samples. Sequences from samples that fell below this threshold were not considered for family-specific downstream analysis.

### 2.4 Genome assembly, annotation and secondary genome structure prediction

Preliminary taxonomic assignment results indicated the presence of two novel near complete genomes from distinct viral families (*Parvoviridae* and *Baculoviridae*). To investigate whether complete genome assemblies could be obtained from the generated data, an extensive set of analysis was performed with the Geneious software v2021.2.2 (Kearse *et al.*, 2012).

For the *Parvoviridae* family, raw reads were mapped to the contig obtained through *de novo* assembly, in order to complete the genome of the virus. The Geneious mapper was used with the highest sensitivity setting. The limits of the new consensus genome sequence were determined through the prediction of the secondary structure of the Inverted Terminal Repeats (ITRs), characteristic of this viral family, using the Vienna package RNAfold tool v. 2.5.1. (Lorenz *et al.* 2011), built in the Geneious software. Both ITR sequences were also aligned using the Muscle software v. 3.8.425 (Edgar, 2004) and had their sequences and secondary structures visually compared to determine their status as homo or hetero-telomeric. Through the EMBOSS Nucleotide Analysis v. 1.1.1 Geneious plugin, the transcription factors associated with the ITR were predicted by the tfscan tool (Rice *et al.*, 2000). The plugin runs the sequence through the TRANSFAC database, finding possible matches set for a minimum of 7 nucleotides of length and 1 nucleotide maximum mismatch. The ITR secondary structure and status are also used to determine genus-level taxonomic status of novel viruses from the *Parvoviridae* family (Cotmore *et al.*, 2019). The open reading frames (ORFs) were identified with the ORFfinder from NCBI with the following parameters: genetic code: 1; start codon: ‘ATG only’; minimal ORF length: 150 nt (https://www.ncbi.nlm.nih.gov/orffinder/).

Multiple contigs related to the *Baculoviridae* family were obtained through *de novo* assembly. All contigs and raw reads were mapped to the most similar reference indicated in BLASTn using Geneious mapper with highest sensitivity. From the mapping results, a consensus genome sequence was obtained. Gene predictions were performed using the FgenesV algorithm for circular genomes available at Softberry, which finds similar proteins in public databases and annotates predicted genes (http://www.softberry.com/berry.phtml). Together with an annotation from the most similar reference also performed on Geneious, final ORFs were defined manually.

### 2.5 Phylogenetic analyses

The assembled viral genomes had their evolutionary history contextualized through similar phylogenetic frameworks. All sequences were aligned with MAFFT v7.505 (Katoh and Standley, 2013) and posteriorly trimmed with TrimAl v.1.4 (Capella-Gutiérrez *et al.*, 2009) with a gap threshold of 0.9, removing sites from alignments composed of 10% or more of gaps. Maximum likelihood trees were inferred with IQ-Tree v.2.1.4 (Minh *et al.*, 2020) with the best fitting model suggested by ModelFinder (Kalyaanamoorthy *et al.*, 2017). Node support was estimated with SH-like approximate likelihood ratio test (SH-aLRT) (Guindon *et al.*, 2010) and ultrafast bootstrap (Minh *et al.*, 2013).

For the *Parvoviridae* genome, the predicted amino acid sequences corresponding to viral structural protein (VP) and non-structural protein (NS1) were identified and had their individual phylogenetic histories inferred with a dataset comprehending the subfamily *Densovirinae*, as described by (Pénzes *et al.*, 2020). For the *Baculoviridae* family, 38 core genes were previously identified through genome sequencing (Javed *et al.*, 2017), as well as an additional set of 25 specific genes in baculoviruses that infect lepidopteran hosts (Garavaglia *et al.*, 2012; van Oers and Vlak, 2007). From those, three were used for phylogenetic analysis. Therefore, nucleotide sequences of *granulin* (*gran*), *late expression factor 8* (*lef-8*) and *late expression factor 9* (*lef-9*) were identified and aligned to previously assembled datasets *(Pénzes *et al.*, 2020; Wennmann et al., 2018).* Once sequences were aligned and trimmed, the genes were concatenated for phylogenetic inference. Original datasets, alignments and phylogenetic trees can be found in **Supplementary file S1**. Additionally, the alignment of concatenated genes was used to infer a pairwise nucleotide distance comparison using the Kimura two-parameter (K2P) model available on MEGA Software v11.0.13 (Tamura *et al.*, 2021).

## 3. Results

### 3.1 Taxonomic composition of the marmoset virome

We characterized the virome from three pools consisting of fecal samples collected from nine free-living hybrids of *C. jacchus x C. penicillata*. The library pool of Afetiva (S_Afe) resulted in 1,740,591 raw reads, which after quality control dropped to 751,600 (43,2%) filtered reads (**Table 2**). A large proportion of the filtered reads had no similarity to any sequences available on Genbank (648,172; 86.24%), 103,428 reads (13.76%) were taxonomically assigned and 690 contigs (192 - 47,091 nt) were *de novo* assembled. The proportion of reads and contigs were 2% and 5% for Viruses, 87% and 89% for Bacteria and 11% and 6% for Eukarya, respectively. Viruses from different taxonomic groups were identified, of which 99% had the closest hits to DNA viruses and 1% to RNA viruses. The viral diversity consisted of several families, including *Siphoviridae*, *Myoviridae* and *Parvoviridae*; unclassified bacteriophages and others with read abundance < 1%. Among the viral families, the *Siphoviridae* family (bacteriophages) was the most abundant, representing 57% of the viruses described.

**Table 2:**
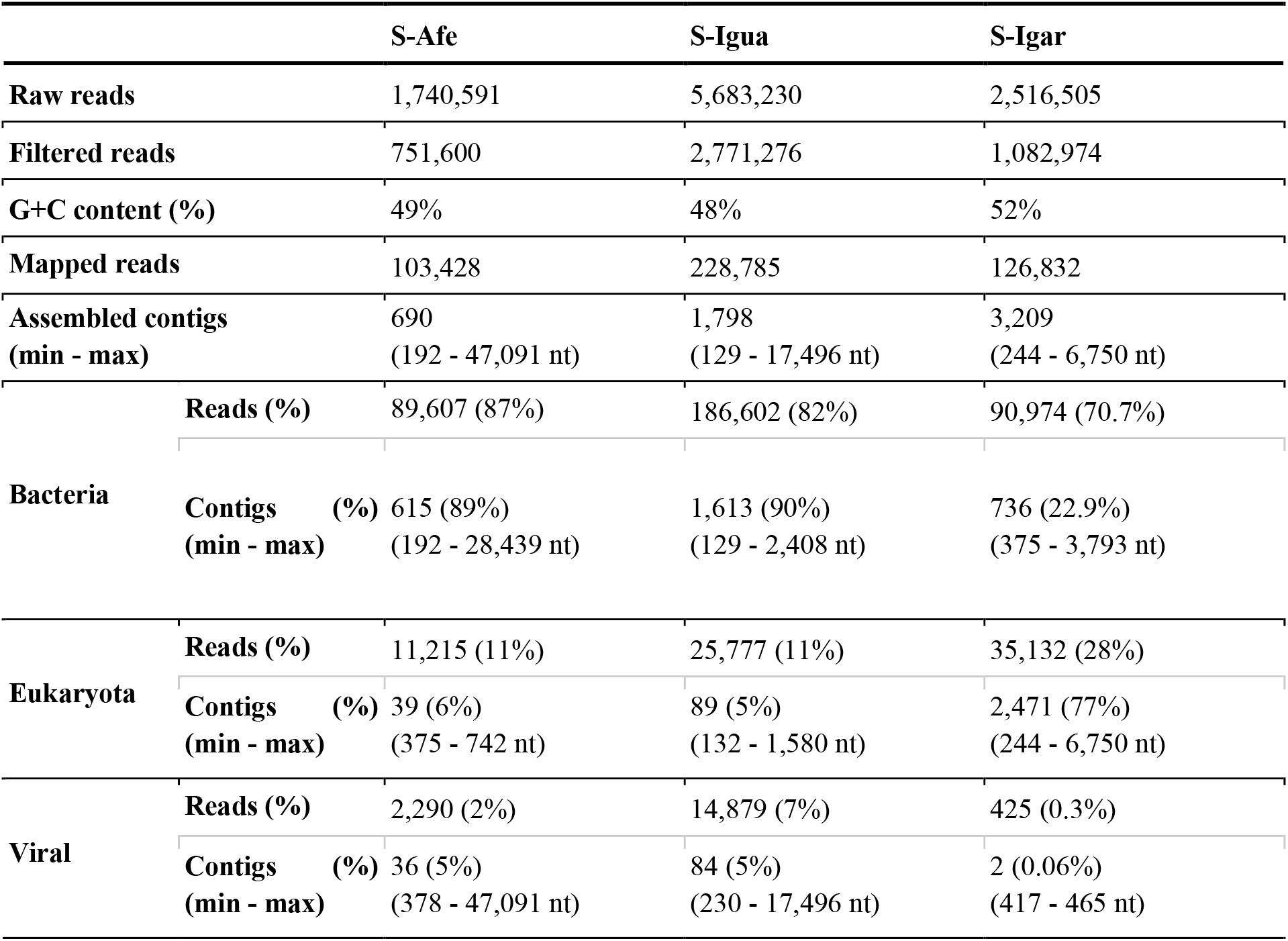
Overview Virome.

The library pool of Iguape (S_Igua) resulted in 5,683,230 raw reads, from which 2,771,276 (48,8%) passed quality control. The proportion of assigned reads was <1%, resulting in 228,785 taxonomic reads and 1,798 assembled contigs (129 - 17,496 nt). The proportion of reads and contigs were 7% and 5% for Viruses, 82% and 90% for Bacteria and 11% and 5% for Eukarya, respectively. Viruses from a wide range of taxonomic groups were identified, of which 98% had the closest hits to DNA viruses and 2% to RNA viruses. The viral diversity was composed by the families *Baculoviridae*, *Myoviridae* and *Siphoviridae*; unclassified *Ribovirus* and others with read abundance < 1%. Among the viral families, the *Baculoviridae* (insect viruses) was the most abundant, representing 90% of the viruses described.

The library pool of Igarapé (S_Igar) resulted in 2,516,505 raw reads (1,082,974 filtered reads; 43%). As in the other libraries, a low proportion of reads had similarity with the sequences available in the Genbank database. A total of 126,832 reads were taxonomically assigned and assembled in 3,209 contigs (244 - 6,750 nt). The proportion of reads and contigs were 0.3% and 0.06% for Viruses, 70.7% and 22.9% for Bacteria and 28% and 77% for Eukarya, respectively. Viruses from diverse taxonomic groups were identified, of which 79% had the closest hits to DNA viruses and 21% to RNA viruses. The viral diversity consisted of families *Siphoviridae*, *Myoviridae, Podoviridae, Autographiviridae, Microviridae, Parvoviridae, Circoviridae, Retroviridae* and *Schitoviridae*; unclassified virus such as *Ribovirus* and *CRESS virus* and others with read abundance < 1%. Among the viral families identified, *Siphoviridae* was the most abundant at 16%. A complete table with viral taxonomic assignments can be found in **Supplementary Table S1**.

### 3.2 Discovery of two novel insect-related viruses

Given trends observed in the abundance of reads and contigs, as well as contig lengths observed for the *Parvoviridae* and *Baculoviridae* families, we were prompted to investigate the occurrence of novel complete genome sequences. In this effort, we characterized two novel viral species and performed comprehensive analyses of their genomes.

#### 3.2.1 Parvovirus

In the library pool S_Afe, a contig of 4,573 nt of length and depth of 431x was detected, pertaining to the *Parvoviride* family with 35,76% identity to nucleotide sequence to *Acheta domesticus mini ambidensovirus* (AdMDV, GenBank: NC_022564.1), detected in common domestic crickets (*Acheta domesticus*) from the United States of America. The final assembly resulted in a genome sequence of 5,309 nt long, with the characteristic *Parvoviridae* 5’ and 3’ ITRs identified. NCBI ORF finder identified the presence of four main ORFs. Three consecutive 5’ ORFs were identified, comprehending 215, 401 and 513 codons, respectively. The second and third ORFs presented long overlapping regions, reflecting distinct reading frames. An additional 3’ ORF of opposite-sense presenting 608 codons was also identified resulting in an ambisense genome (ssDNA (+/−)) (**Figure 1A**).

**Figure 1:**
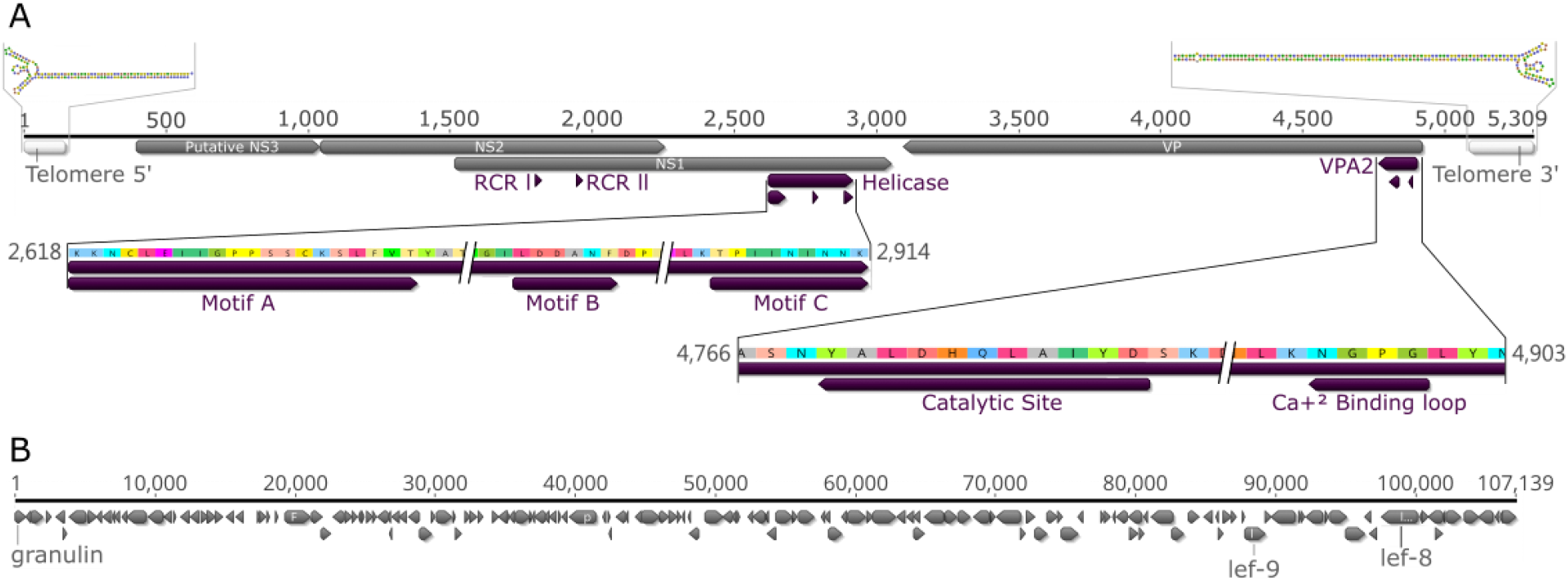
Genome characterization of the novel Parvovirus and Baculovirus identified in hybrids of *Callithrix sp.* feces by viral metagenomics. ORFs and transcription direction are indicated as arrows. A) Complete genome of *Fecalis tritonambidensovirus 1* (TfDV). Main genes are indicated as gray arrows, motifs are represented as purple arrows and telomere region structure is represented within white boxes. The predicted structures of the ITRs are above its respective region. B) Complete genome of *Callithrix fecalis granulovirus* with its 153 ORFs indicated as gray arrows. Named genes correspond to those commonly used for viral family phylogenetic analysis and used in the present work.

BLASTp tool revealed that the 512-aa protein had 31.8% identity with 89% coverage (E-value 6e-64) with the NS1 and the 607-aa protein had 27.8% identity and 91% coverage (E-value 1e-35) with the VP, both related to AdMDV (Accession No: YP_008658568.1 and YP_008658570.1 respectively). Interestingly, the 400-aa protein was closest to the NS2 protein of *Phoenicurus auroreus ambidensovirus* (Accession No: QTE03811), found in feces of Daurian redstart (*Phoenicurus auroreus*) in Asia (identity 27.45%, coverage 60% and E-value 3e-09). The smallest protein of 214-aa is a putative NS3 protein from *Densovirinae*, with no similarity to protein sequences available in the database. The NS3 is under the control of the NS3 5’ promoter (pNS3) and the promoters for NS2 (pNS2) and NS1 (pNS1) are overlapping and upstream of the NS2 initial codon, while VP is under the control of the pVP promoter. Three core promoter elements (CPEs) - including the B recognition element (BRE), TATA box and initiator element (Inr) - and the downstream promoter element (DPE) are in the pNS3, pNS1/NS2 and pVP promoter regions, whereas the MTE (motif ten element) was not found.

Based on sequence comparison to known densoviruses, two replication initiator motifs called rolling circle replication I and II (RCR I and RCR II) and three helicase motifs were identified in NS1 (**Figure 1A**). The RCR I is located at position 96-103 aa (_96_HxHxxHDC_103_) containing two conserved histidine residues (H96 and H98) for metal binding. The RCR II is located at position 145-151 aa (_145_Yxxxxxx_151_) and contains a tyrosine residue (Y145) important to participate in the cleavage-linkage reaction. The NS1 also contains the DNA-dependent ATPase/helicase domain with the Walker A, B, and C motifs (aa 369–467). The viral phospholipase A_2_ (vPLA_2_) was identified in the N-terminal extension of VP protein (aa 6-51) with the calcium binding loop (_12_GPGN_15_) and catalytic site (_28_DxxAxxHDxxY_38_) found in all *Densovirinae* subfamily members. Sequence alignment of the putative NS3 protein revealed four conserved cysteine residues (_113_CxxC55xCxxC_175_) comprising the conserved Zn-finger motif (apoptosis inhibitors) present in all NS3-like arthropod virus proteins. The ORFs and motifs can be found in **Table 3**.

**Table 3:**
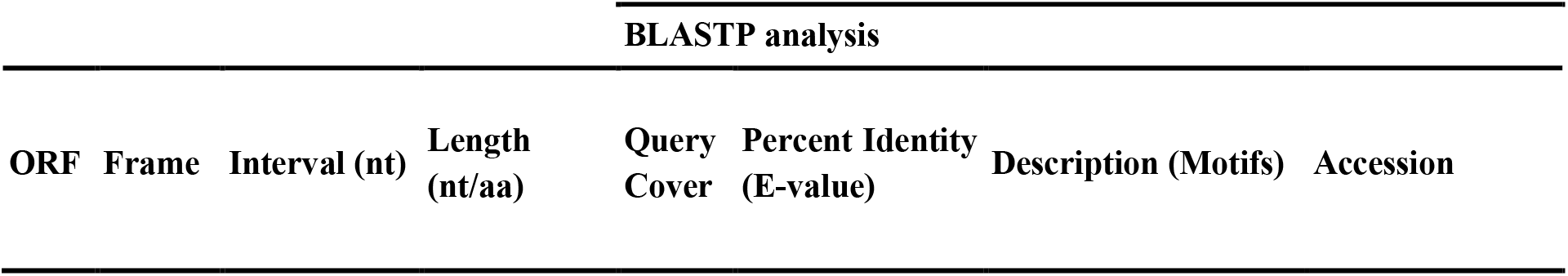
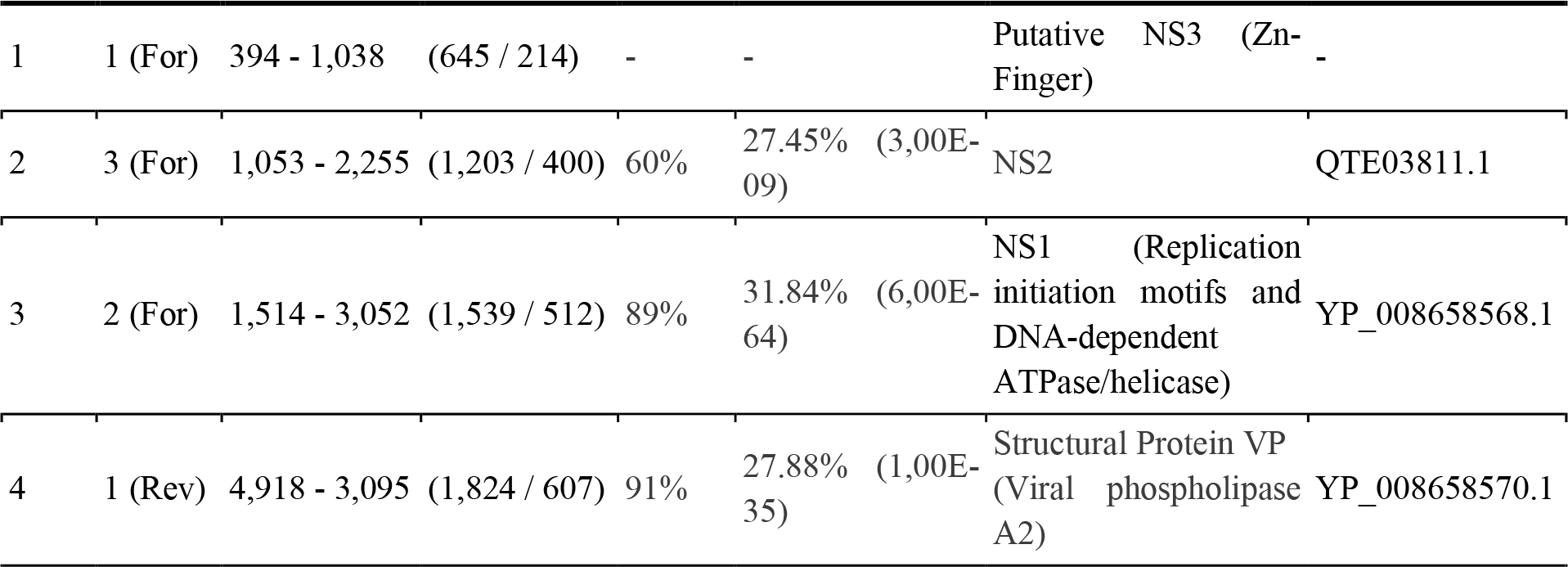
ORFs and motifs.

The 5’ and 3’ telomeric ends of the genome of the new densovirus are, respectively, 145 and 236 nt and form ITRs. The ITRs form imperfect palindromes that conform into secondary structures and present higher GC content (5’ ITR: 62.8% and 3’ ITR: 52.3%) than the genome as a whole (36.9%). Using Geneious, we identified the minimum free-energy secondary structure of the newly described densovirus (**Figure 1A**). Both ends develop similar trident-shaped secondary structures, and when aligned (data not shown), the shorter 5’ ITR is complementary to the central portion of the 3’ ITR (trident loop and beginning of stem). This classifies the novel *Densovirinae* herein described as homotelomeric. The most common structure found in other genera of this subfamily are J-shaped telomeres.

The NS1 phylogenetic tree reveals the new densovirus as a sister group to the single representative of the *Miniambidensovirus* genus (AdMDV) with robust node support (SH-aLRT = 100; UFBoot = 99) (**Figure 2**). The described FtDV represents a novel genus, according to the ICTV genera demarcation criteria to the *Parvoviridae* family (*i.e.*, < 40% NS1 amino acid sequence identity) (Walker *et al.*, 2021; Penzes *et al.*, 2020). Different from NS1, the putative VP protein clustered within the recently described sequence (*Muscodensovirus*) with limited branch support (SH-aLRT = 68; UFBoot = 61.3) (**Supplementary Figure S1**). Thus, due to an ambisense genome, format of trident ITRs and the sample type, we named the new virus provisionally as *Fecalis tritonambidensovirus* 1 (FtDV) according to the new binomial nomenclature stipulated by ICTV.

**Figure 2:**
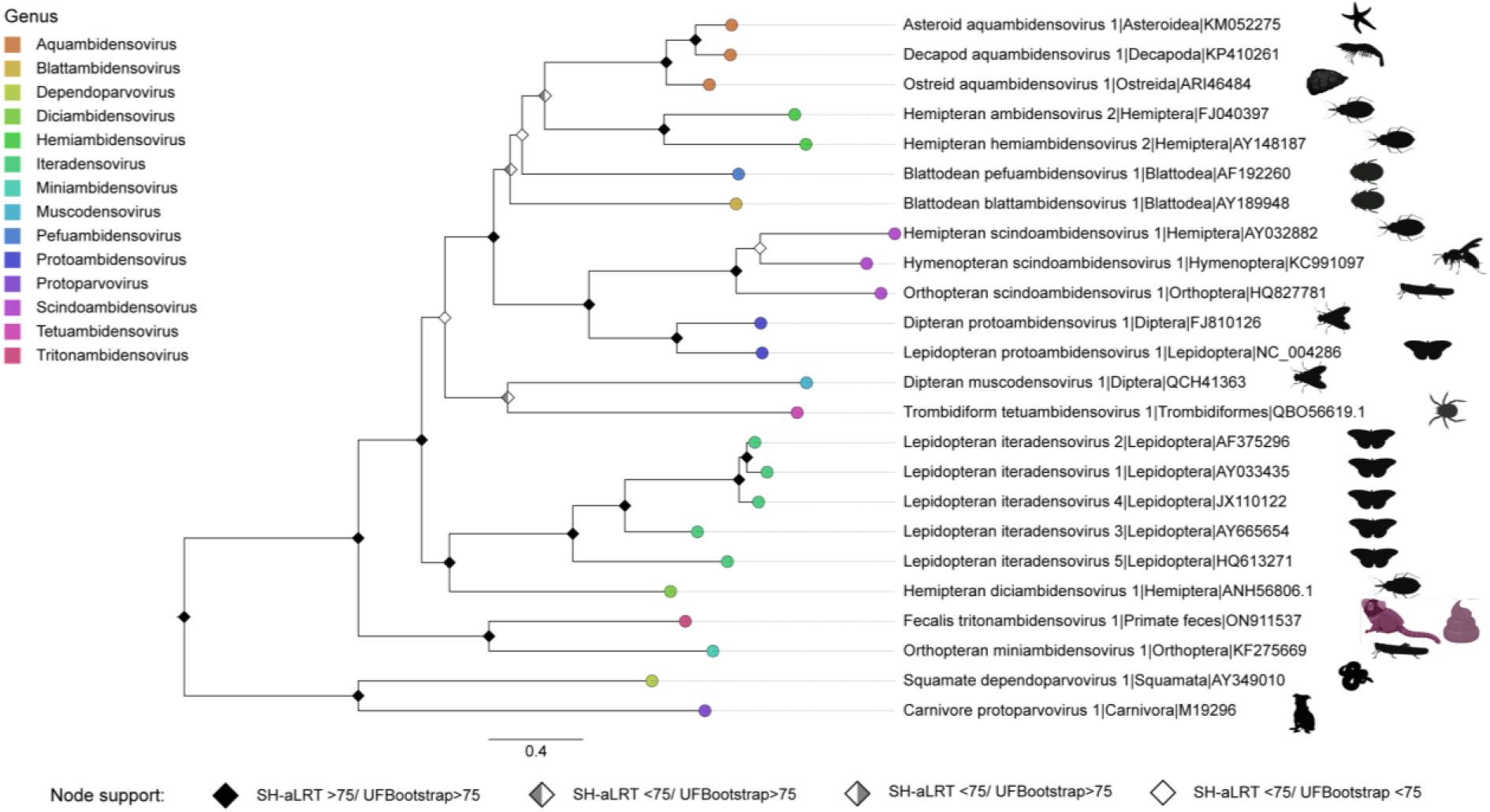
Maximum likelihood phylogeny of the *Densovirinae* subfamily, *Parvoviridae*. Phylogeny of the NS1 gene for the *Densovirinae* subfamily, 416 amino acids length under LG+F+I+G4 model. *Fecalis tritonambidensovirus 1* comprehends a new genus, sister group to the AMDv. Tip labels are colored according to the Parvoviridae genus used in the dataset, the proposed genus *Fecalis tritonambidensovirus 1* tip is colored in fuchsia. Viral hosts described in literature had their order silhouette represented in black. *Fecalis tritonambidensovirus 1* (TfDV) obtained in the study has its host colored in fuchsia. Node labels colored in black represent support for SH-aLRT and Bootstrap equal or superior to 75. When only one of the parameters was superior it is represented in gray, while the other in white.

#### 3.2.2 Baculovirus

From a *de novo* assembly using reads from the S_Igua pool, a total of 23 *Baculoviridae* contigs were identified, ranging from 230 to 17,496 nt. Following a megablastn analysis, this contig showed 85.7% similarity with *Clostera anastomosis granulovirus B* (ClasGV-B; GenBank: NC_038371.1), a virus of the *Betabaculovirus* genus, specific to the Lepidoptera insect order (moths and butterflies). All 23 contigs were then mapped to this reference genome, covering 88.6% of its extension. After mapping raw reads to this preliminary assembly, genome coverage raised to 100% with 60.8% similarity. The resulting consensus genome sequence had 107,139 nt, a GC content of 34.4% and complementary ends that confirmed circularization.

A total of 153 ORFs with 140 nt or more in length were identified (**Figure 1B**). As previously established, nt 1 was defined based on the adenine of the *granulin* gene, which also defines orientation of the circular genome. That said, 80 genes were clockwise directed and 73 were counterclockwise. From the 38 core genes, 26 were also identified complete in this novel virus, including two genes necessary for phylogenetic analyses: *orf255* (*lef-8*) and *orf155* (*lef-9*), comprehending 875 and 499 codons, respectively. The other 12 genes were found either incomplete (n=7) or truncated (n=4), and one was not identified at all. Additionally, 20 of the 25 genes found in lepidopteran baculoviruses were present in the genome characterized, including the *granulin* gene (*orf1*; 249 codons), also necessary for phylogeny. Finally, 13 ORFs showed no homology with other genes, being, so far, unique to this virus.

Phylogenetic analysis based on *granulin*, *lef-8* and *lef-9* genes revealed that the novel virus clusters into the *Betabaculovirus* clade, supporting its close relation to ClasGV-B (**Figure 3**). In addition, for lepidopteran-specific baculoviruses, inferences of nucleotide pairwise distances with Kimura two-parameter (K2P) have also been validated for species demarcation: distances up to 0.021 designate viruses belonging to the same species, values up to 0.072 relies on more information and over this threshold indicates viruses of different species *(Wennmann et al., 2018).* In this case, the genetic distance from the closest reference (ClasGV-B) was calculated as 0.144, confirming that the virus proposed here belongs to a new species, provisionally named *Callithrix fecalis granulovirus* (CafeGV).

**Figure 3:**
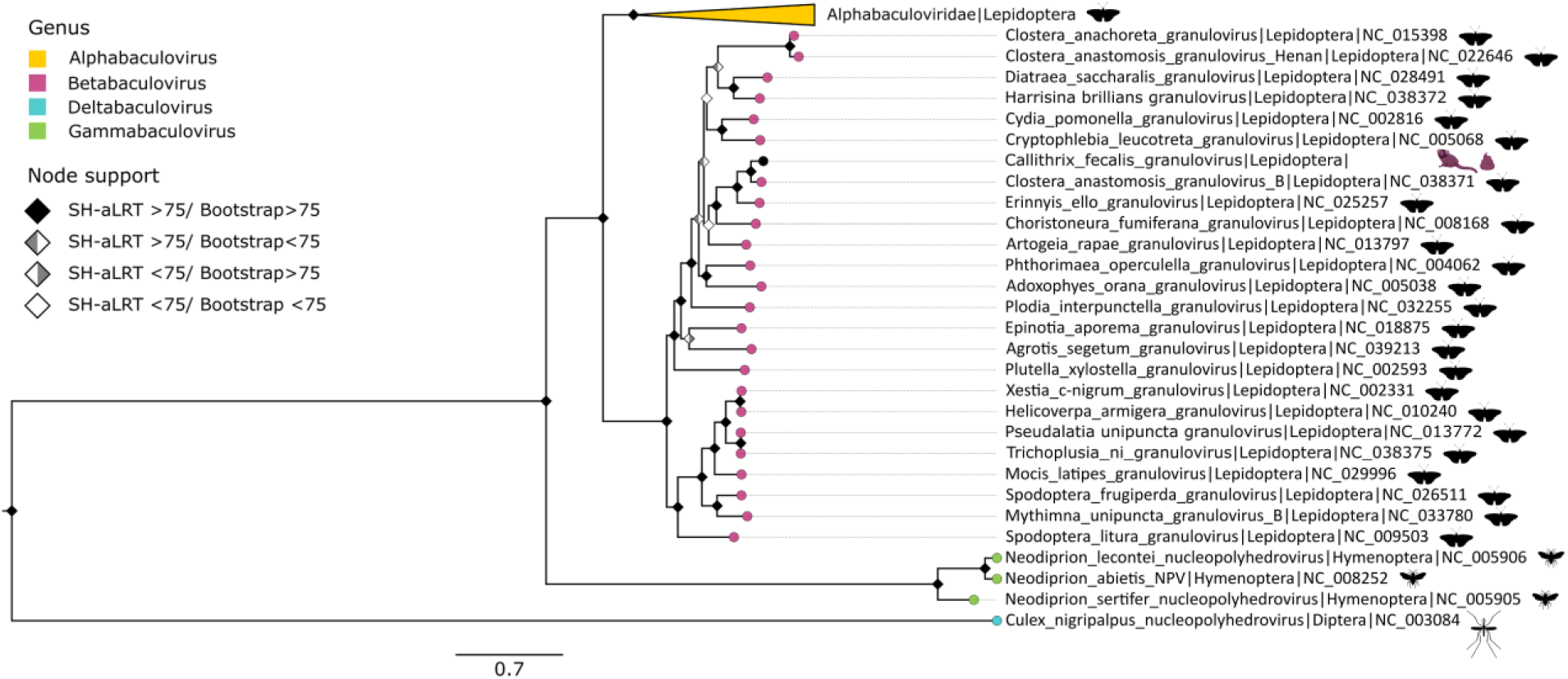
Maximum likelihood phylogeny of the *Baculoviridae* family. Nucleotide phylogeny of *granulin* (*gran*), *late expression factor 8* (*lef-8*) and *late expression factor 9* (*lef-9*) core genes concatenated, 4.579 bp length under SYM+I+G4 model. The baculovirus identified in the study belongs to the previously described *Betabaculovirus* genus, sister to *Clostera anastomosis granulovirus B*. Circular tip labels are colored according to the *Baculoviridae* genus used in the dataset - *Alphabaculovirus* genus is collapsed and colored in yellow, *Betabaculovirus* in pink, *Deltabaculovirus* in blue and *Gammabaculovirus* in green. *Callithrix fecalis granulovirus* obtained in this study is evidenced by a black tip. Node labels colored in black represent support for SH-aLRT and Bootstrap equal or superior to 75. When only one of the parameters was superior, it is represented in gray, while the other is in white.

### 3.3 Arthropods signals in metagenomes

In all pools, sequences related to arthropods were obtained. These sequences were often related to groups of hosts known for viruses related to the ones described in this study. In the S_Afe library, where FtDV was found, about 8% (948) of Eukarya reads were mapped to the arthropod phylum, mounted in 25 contigs (375 to 742 nt in length) (**Supplementary Table 2**). The main families found in terms of relative abundance of contigs were 48% *Pyroglyphidae*, 12% *Sarcoptidae* and 4% *Psoroptidae* (Order Sarcoptiformes - mites); 20% of *Tephritidae* (Order Diptera - fruit fly), followed by 4% each of *Pyralidae* (Order Lepidoptera - moths), *Araneidae* (Order Araneae - spiders), *Ixodidae* (Order Ixodida - tick) and *Trombiculidae* (Order Trombidiformes - mites) (**Figure 4**).

**Figure 4:**
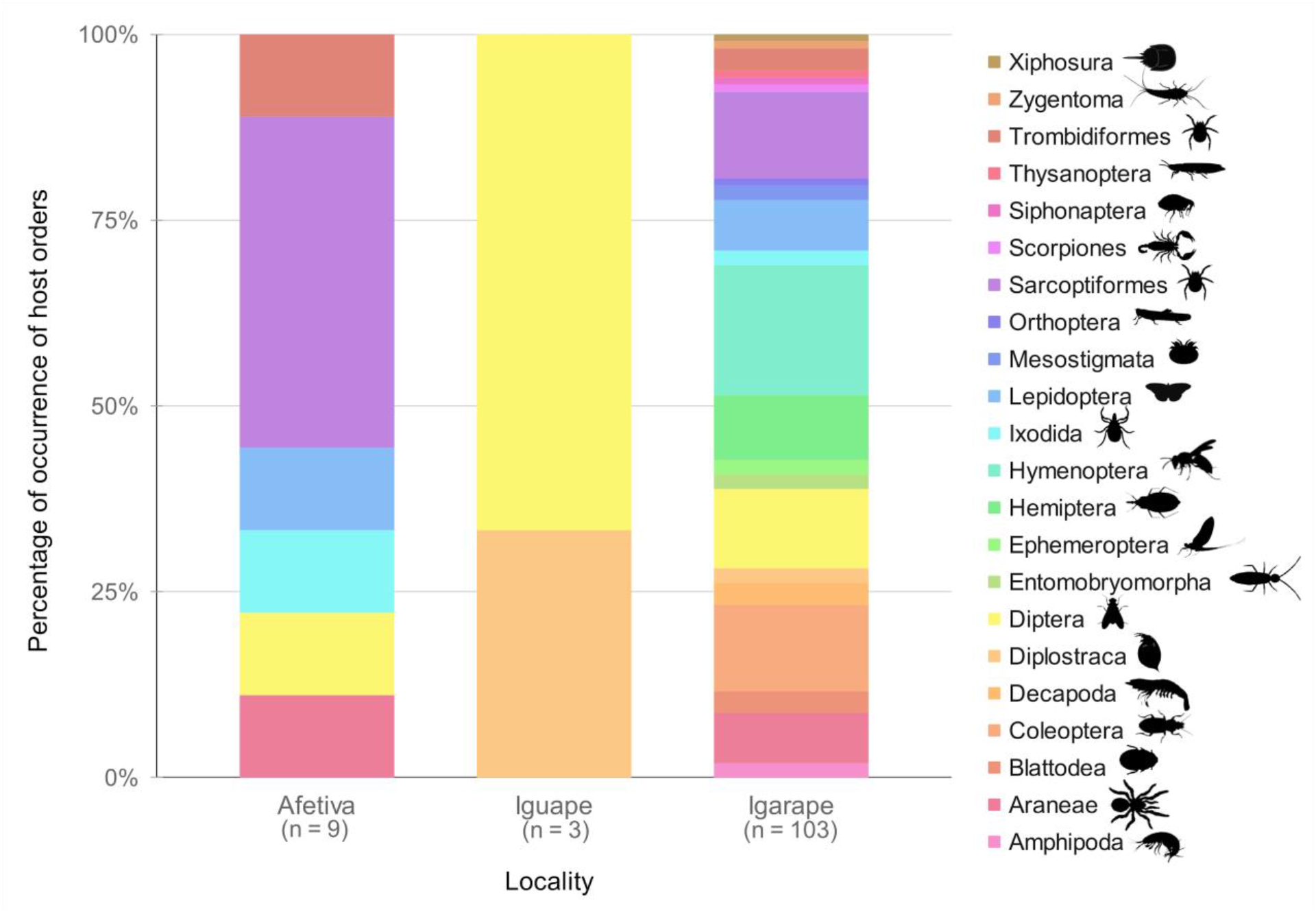
Relative abundance of host order identified by contig taxonomical assignment. Colors and legend on the right refer to the different host orders identified by locality. Columns represent the percentage of host families within each identified order. A total of 9 families in 6 different orders were identified in Afetiva, 3 families in 2 different orders in Iguape and 103 families in 22 different orders in Igarape.

In the Iguape pool (S_Igua), where CafeGV was sequenced, about 1% (307) of Eukarya reads were mapped to the arthropod phylum. However, only three contigs were obtained (412, 413 and 1131 nt in length). The arthropod families corresponding to these contigs were 33.3% each for *Calliphoridae*, *Tephritidae* (both Diptera order - fruit fly) and *Daphniidae* (Diplostraca order - water flea).

Although no arthropod-related viruses were found in the Igarape pool (S_Igar), this library had the greatest amount and diversity of arthropod sequences. About 78% (27,315) of Eukarya reads were mapped to the arthropod phylum, mounted in 2,372 contigs (244 to 6,750 nt in length). In all, 21 orders encompassing 96 families of arthropods were sequenced, where the most abundant were orders Sarcoptiformes, mainly the families *Pyroglyphidae* (55%) and *Sarcoptidae* (17,6%), and Diptera, most abundant family *Tephritidae* (17,2%).

## 4. Discussion

High throughput sequencing is a powerful tool for microbial metagenomic analysis. It has been used to discover a large diversity of novel viruses that could not have been identified by traditional methods (Cosentino *et al.*, 2022; D’arc *et al.*, 2020; Harvey and Holmes, 2022; Shi *et al.*, 2016; Zayed *et al.*, 2022). Following previous virome studies based on non-invasive samples (A Duarte *et al.*, 2019; Bodewes *et al.*, 2014; Wang *et al.*, 2019), we characterized the viral metagenomic of fecal samples of *C. jacchus x C. penicillata* hybrids, the first virome analysis from introduced free-living marmosets. Consistent with the observation that it is particularly challenging to identify virus-host associations from fecal samples using HTS *(Ge et al., 2012; Waller et al., 2022; Wang et al., 2019)*, we identified two new likely dietary-related viruses from the *Parvoviridae* and *Baculoviridae* families, both related to insect hosts.

Although the aim in virome studies is sequencing viruses, a large proportion of non-viral genomes were also sequenced in the present study. Previously projects have shown through metagenomics in feces the great diversity of insects consumed by NP, revealing an existing ecological network (Pickett *et al.*, 2012). Silvestre *et al.* (2016) indicated that arthropods constitute about 61% of the diet of the *C. jacchus*. Therefore, it is conceivable to assume that the abundance of insect-related viruses in our samples is a reflection of the marmoset insectivorous diet (Pickett *et al.*, 2012). Noticeably, most of the novel sequences related to insect viruses have low genetic identities with known viruses, indicating that a number of novel viruses exist in the largely uncharacterized virome of Neotropical insect populations (Ali *et al.*, 2021; Maia *et al.*, 2019; Scarpassa *et al.*, 2019). In addition, the virome of marmosets’ feces was very similar to that observed in the oral cavity of free-living capuchin monkeys (*Sapajus nigritus*) *(Dos Santos et al., 2020).* Both primates are considered opportunistic omnivores and feed on a wide variety of food types, including fish, crustaceans, and insects (Galetti and Pedroni, 1994). Insect-related viral families were also sequenced in capuchin monkeys, demonstrating how metagenomics contributes to the characterization of viruses related to the hosts’ diets (D’arc *et al.*, 2018; French *et al.*, 2022; Geoghegan *et al.*, 2021).

We report the complete genome sequence of a novel viral species from the *Parvoviridae* family. This family comprises non-enveloped viruses with linear ssDNA genomes (3.7–6.3 Kb) and characteristic terminal hairpins (Cotmore *et al.*, 2014). Although the putative NS3 had no similarity to any Genbank sequence, analysis of the amino acids sequence exhibits a conserved Zn-finger motif from NS3-like of several insect viruses (Krupovic and Koonin, 2014). ITRs are also essential for replication, encapsulation and genome integration (Siegl and Tratschin, 1987; Zádori *et al.*, 2001). Due to the presence of palindromic repeats, the most common structure found in the *Densovirinae* subfamily are J-shaped telomeres, but they can also be found in T-, I-, Y-, U- or simple hairpin-like structures (Laugel *et al.*, 2022). However, the predicted structure of the ITRs for this new virus presented a unique trident-like shape. Following ICTV’s genus demarcation (Pénzes *et al.*, 2020), the new species belong to a new genus. Thus, due to the characteristic of the trident shaped ITRs, the ambisense genome architecture and the type of sample, we propose to name it as *Fecalis tritonambidensovirus 1* (FtDV). Interestingly, in the library in which the new FtDV was identified (S_Afe), only Diptera and Lepidoptera orders are known hosts for densoviruses (genus *Protoambidensovirus*) *(Pénzes et al., 2020)* and both orders were retrieved from our data. However, FtDV is more closely related to an Orthoptera virus which was not found in the library.

We were also able to characterize the complete genome sequence of a novel member of the *Baculoviridae* family. Baculoviruses are insect-specific pathogens, composed of proteic occlusion bodies (OBs) containing enveloped viruses with circular dsDNA genomes, ranging from 80 to 180 kbp (Harrison *et al.*, 2018). They were traditionally classified based on host range, which include orders Lepidoptera, Hymenoptera and Diptera (Herniou *et al.*, 2003), and the OB’s composition, which may be polyhedral or ovocylindrical. However, recent phylogenetic approaches demonstrated that phylogenies using highly conserved genes are also a reliable method of classification (Lange *et al.*, 2004; Wennmann *et al.*, 2018). The ICTV guidelines for new virus names indicate a pattern of host specific name, followed by the designation of OB’s composition ((Harrison *et al.*, 2018). Since it is not possible to define the direct host of this novel virus and considering we used feces samples of *Callithrix sp.*, we provisionally named it *Callithrix fecalis granulovirus* (CafeGV). The phylogenetic relationship was done based on three core genes, as accepted in the literature ((Lange *et al.*, 2004; Wennmann *et al.*, 2018)). We note that a number of putative core genes are incomplete or absent in the new genome and highlight this new virus may display more genomic plasticity than previously appreciated for the genus. Additional experiments are mandatory to further clarify its genome structure. Finally, in the S_Igua library, where this new *Betaculovirus* was identified, only the Diptera and Diplostraca (Crustacea) orders were sequenced, while the Lepidoptera order, host of all previously described *Betabaculoviruses*, was not detected (Jehle *et al.*, 2006). However, it is possible that the original host was underrepresented in this sample or could not be assigned to the right taxonomic level.

It is important to highlight that the arthropod-related sequences obtained by HTS in *Callithrix sp.* feces may not necessarily reflect the diversity of arthropods in the diet, since viral metagenomics has a compositional bias toward sequencing viral agents (Kumar *et al.*, 2018). Thus, it becomes difficult to clearly determine whether the likely hosts have been disadvantaged in the HTS or the range of hosts for the mentioned viral groups is wider than previously known. To untangle both scenarios, further investigations into the diversity and evolution of insect viruses are needed.

## 5. Conclusion

Our study provides, for the first time, an overview of the fecal virome of introduced free-living *Callithrix sp.* hybrids. Through viral metagenomic sequencing, it was possible to characterize two novel dietary-related complete viral genomes, culminating in the description of a new provisional genus for the subfamily *Densovirinae* and a new viral species for the genus *Betabaculovirus*, contributing to the expansion of the families *Parvoviridae* and *Baculoviridae*, respectively. Further studies are needed to improve our knowledge of the virome of the marmoset hybrids and the parental species, *C. jacchus and C. penicillata*, allowing us to better understand the circulation of viruses among these animals and their impact on endemic primate species.

## Data Availability

Sequences generated in the present work were submitted to Genbank under the accession numbers: ON911537 and OP589943. The metagenomic data set is also available in the NCBI SRA repository under Bioproject PRJNA847605 (SAMN28946322 - SAMN28946324).

## Conflict of interest

The authors declare no conflicts of financial or personal interests.

## Funding

This work was supported by the Conselho Nacional de Desenvolvimento Científico e Tecnológico/CNPq (grant number 313005/2020-6 to AFS) and Fundação de Amparo à Pesquisa do Estado do Rio de Janeiro/FAPERJ (grants numbers E26/202.738/2018, E26/210.122/2018, and E26/211.040/2019 to AFS).

## Acknowledgements

We thank the field team of the *Associação do Mico-Leão-Dourado* (AMLD) for their help in collecting samples and logistic support in the field. We also thank the members of the LDDV laboratory for their fantastic teamwork.

## Notes

### Competing Interest Statement

The authors have declared no competing interest.

