## Supplementary figures and images for "Metagenomic analysis reveals novel dietary-related viruses in the gut virome of marmosets hybrids (*Callithrix jacchus x Callithrix penicillata*), Brazil"

### Supplementary Figure 1

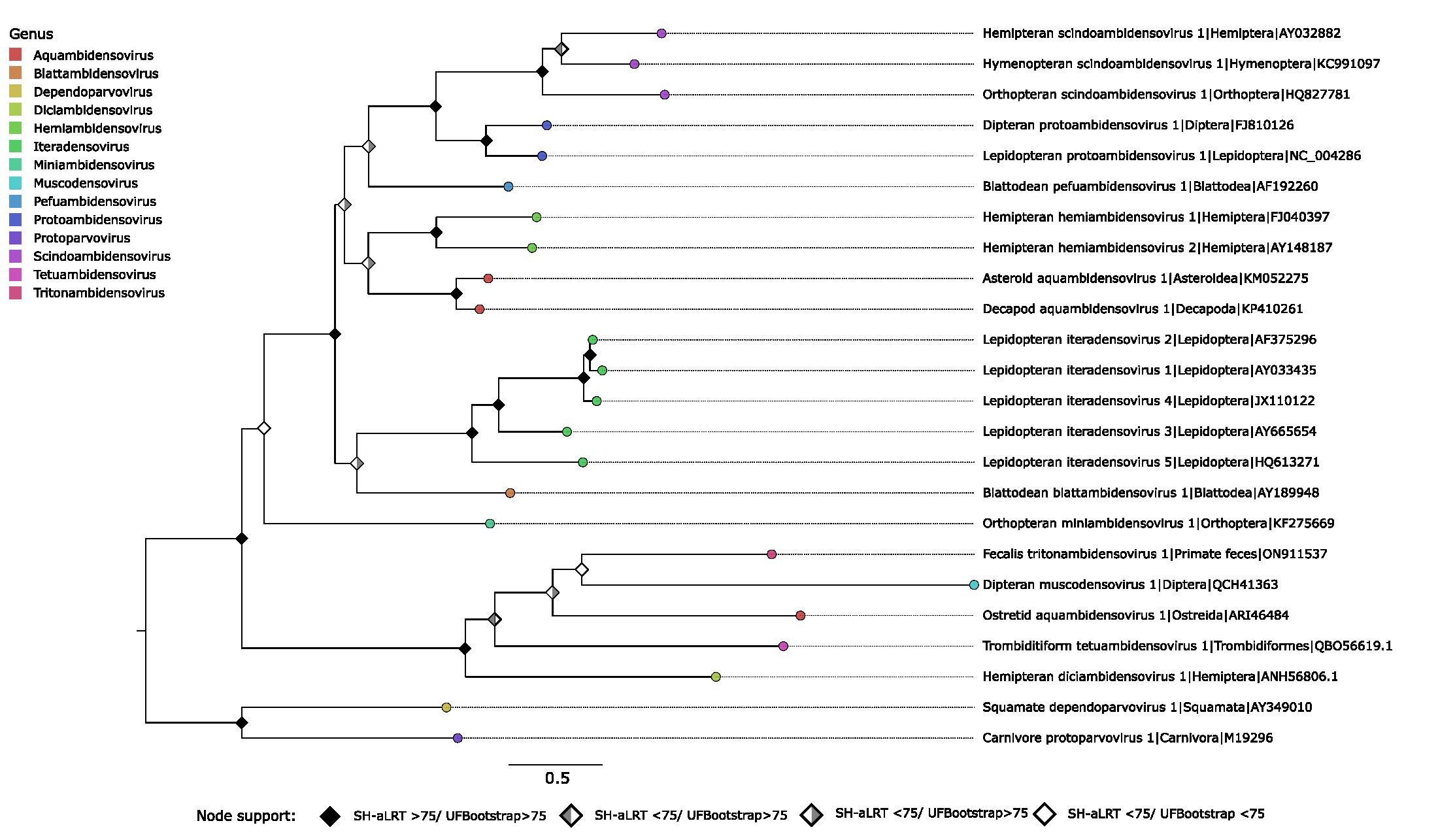
